# Sexually dimorphic influence of the circadian clock gene *Bmal1* in the striatum on alcohol intake

**DOI:** 10.1101/2020.09.16.299842

**Authors:** Nuria de Zavalia, Konrad Schoettner, Jory A. Goldsmith, Pavel Solis, Sarah Ferraro, Gabrielle Parent, Shimon Amir

## Abstract

The gene *Bmal1* (brain and muscle Arnt-like 1) plays an obligatory role in the generation of circadian rhythms in the suprachiasmatic nucleus (SCN), the master circadian clock in mammals [1–5]. Notably, *Bmal1* is widely expressed in mammalian brain [6], and perturbations in *Bmal1* expression in select forebrain regions cause behavioral disturbances that are independent of the SCN, such as disturbances in sleep architecture, and in cognitive and affective behaviors [1, 7–15]. Interestingly, gene association studies in humans and in animals suggest that *Bmal1* may influence the propensity to consume alcohol, and that polymorphisms in *Bmal1* may confer risk for alcohol dependence and related disorders [16–20]. However, research has not yet provided evidence of a causal role of *Bmal1* in the control of alcohol intake. We investigated voluntary alcohol consumption in conditional knockout mice that lack *Bmal1* exclusively in the striatum, which is an important structure in the control of alcohol intake and preference [21–26]. Experiments were carried out in both male and female mice in order to account for the known sex differences in alcohol consumption [27–31] and in striatal functioning [32–36], as well as in the expression of clock genes and in the impact of circadian clocks on behavior [37–44]. We found that, in both males and females, selective deletion of *Bmal1* from principal medium spiny neurons (MSNs) of the striatum significantly altered voluntary alcohol intake and preference. Strikingly, the effect of *Bmal1* deletion was sexually dimorphic. Whereas in males, deletion of *Bmal1* augmented alcohol intake and preference, in females, the same deletion suppressed alcohol intake and preference. Interestingly, striatal deletion of the clock gene *Per2,* which interacts with *Bmal1* in the generation of circadian rhythms [4], and which has been shown to affect alcohol consumption in male mice [45], mimicked the effect of *Bmal1* deletion, albeit only in males. These results show that *Bmal1* in MSNs of the striatum exerts a sexually dimorphic influence on alcohol intake in mice, moderating intake in males, possibly via *Per2,* and promoting heightened intake in females, independently of *Per2*. We propose that a sexually dimorphic mechanism in the function of *Bmal1* in the striatum contributes to sex differences in the propensity to consume alcohol in mice. Whether such mechanism contributes to sex differences in other striatum-dependent appetitive and consummatory behaviors remains to be investigated.

## Expression of BMAL1 in Striatal MSNs and Generation of Striatal *Bmal1* Knockout Mice

Striatal MSNs play an important role in alcohol neuroadaptation and alcohol intake and preference [22, 46–48]. They constitute approximately 95% of the entire striatal neuronal population in mice [49, 50] and consist of two intermixed populations, one that projects directly (striatonigral), and one that projects indirectly (striatopallidal) to the output nuclei of the basal ganglia [51]. Striatonigral MSNs are distinguished by the expression of D1 dopamine (DA) receptors, and striatopallidal MSNs by the expression of D2 DA receptors [52]. Both MSNs subtypes express Gpr88, a striatum-specific G-protein coupled receptor [53–55]. Using fluorescence immunohistochemistry in tdTomato-D1/GFP-D2 double transgenic reporter mice [56], we confirmed that both D1 and D2 DA receptor bearing neurons in the striatum express BMAL1 (Figure 1A). Moreover, using transgenic mice that express Cre and green fluorescent protein (GFP) under control of the *Gpr88* promoter, we confirmed that BMAL1 is expressed in Gpr88-bearing neurons of the striatum (Figure 1B).

**Figure 1.**
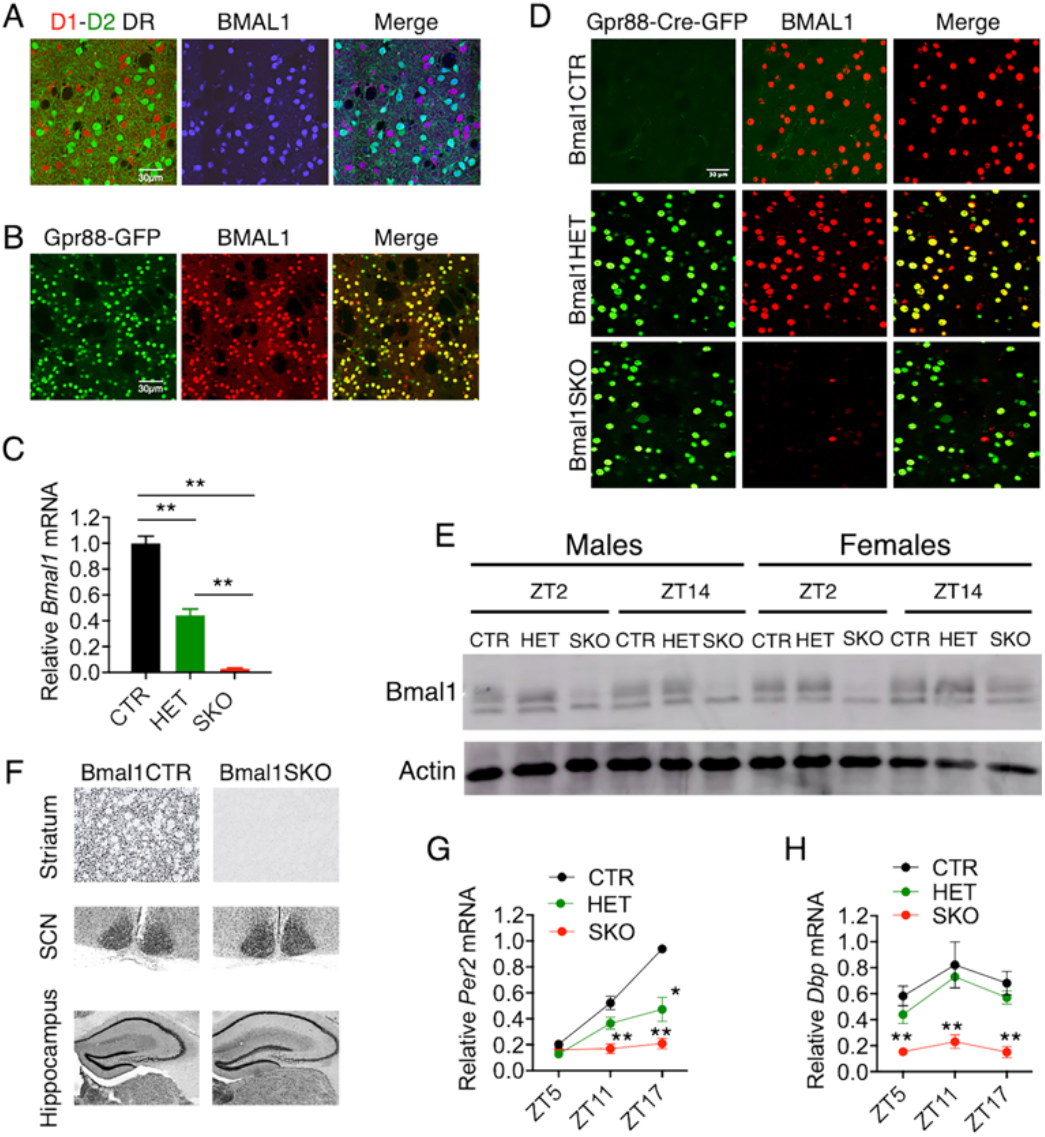
BMAL1 expression is dampened in MSNs of conditional *Bmal1* knockout mice. **A)** Representative images of immunofluorescence staining in tdTomato-D1/GFP-D2 double transgenic reporter mice showing that BMAL1 (blue) is expressed in D1 (red) and D2 (green) dopamine receptor bearing neurons of the dorsal striatum (scale bar = 30 μm). **B)** Representative images of immunofluorescence staining showing that BMAL1 (red) is expressed in Gpr88-Cre-GFP (green) positive MSNs of the mouse dorsal striatum (scale bar = 30 μm). **C)** Quantitative PCR analysis of dorsal striatal tissue shows a clear effect of gene dosage on *Bmal1* mRNA levels, with *Bmal1* knockout mice displaying almost no expression, and *Bmal1* heterozygote mice exhibiting approximately half of the mRNA levels as control mice in the striatum. There were significant differences between all groups (**P<0.0001, unpaired two-tailed t test). **D)** Representative image of immunofluorescence staining showing that *Bmal1* knockout mice which express Cre-GFP in dorsal striatal MSNs, and have two floxed *Bmal1* alleles, display depleted BMAL1 in GFP-positive cells. *Bmal1* heterozygote mice which express Cre, but only possess a single floxed *Bmal1* allele, retain BMAL1 expression. Control mice which contain two floxed *Bmal1* alleles, but lack Cre, retain BMAL1 expression. **E) A** representative western blot image confirming BMAL1 depletion in the dorsal striatum of SKO male and female mice at two different times of the day. **F)** Representative image of immunohistochemistry staining showing expression of BMAL1 in the hippocampus and SCN, but not in the striatum of *Bmal1* knockout mice. **G)** Quantitative PCR analysis showing that *Per2* mRNA levels were downregulated and its rhythm was blunted across the light-dark cycle in the dorsal striatum of conditional *Bmal1* knockout mice. (*P<0.01, **P<0.005, unpaired two-tailed t test). **H)** Quantitative PCR analysis of Dbp mRNA expression shows a downregulation of this gene in the dorsal striatum of conditional *Bmal1* knockout mice. (** represent significant difference from control, p<0.005, unpaired two-tailed t test).

Mice with specific deletion of *Bmal1* from MSNs of the striatum were generated using the Cre-*lox* recombination strategy. C57BJ/6 mice homozygous for floxed alleles of the *Bmal1* locus ([Bmal1fl/fl], JAX, stock number 7668) were crossed with transgenic mice that express Cre recombinase and GFP under control of the *Gpr88* promoter ([Gpr88Cre/+], JAX, stock number 22510) to yield striatal specific *Bmal1* knockout mice (Gpr88Cre/+; Bmal1fl/fl [Bmal1SKO]), as well as heterozygotes (Gpr88Cre/+; Bmal1fl/+ [Bmal1HET]) and wild type controls (Gpr88+/+; Bmal1fl/fl [Bmal1CTR]). The resulting Bmal1SKO and Bmal1HET male and female mice were similar to their littermate controls in weight. Quantitative real-time polymerase chain reaction (qPCR) analysis of striatal tissue punches (n = 3/genotype) revealed a significant reduction of mRNA levels within the floxed *Bmal1* locus to less than 5% of control levels in Bmal1SKO mice (P<0.0001, unpaired two-tailed t test) and less than 50% of control levels in Bmal1HET mice (P<0.0001, unpaired two-tailed t test) (Figure 1C). Using fluorescence immunohistochemistry, we confirmed that striatal brain sections from Bmal1CTR and Bmal1HET mice expressed BMAL1, whereas striatal sections from Bmal1SKO mice lacked BMAL1 immunostaining (Figure 1D). The reduction of BMAL1 protein in the striatum was further confirmed by western blotting analysis of striatal tissue punches. In both male and female Bmal1SKO mice, the expression of BMAL1 was substantially lower at zeitgeber time (ZT) 2 and ZT14 compared to control animals (Figure 1E). Moreover, we confirmed that the deletion of *Bmal1* is restricted to the striatum by showing presence of BMAL1 staining in the SCN and hippocampus of Bmal1SKO mice (Figure 1F). To determine whether deletion of *Bmal1* disrupted the striatal circadian clock we studied the expression of the clock gene, *Per2*, which depends on *Bmal1* for expression and daily cycling (n=3/timepoint). In Bmal1CTR mice, *Per2* mRNA levels in the striatum peaked at night (ZT17). In contrast, in Bmal1SKO mice, the levels of *Per2* mRNA were significantly downregulated at ZT11 (CTR versus SKO, P<0.005, unpaired two-tailed t test) and ZT17 (P<0.005, unpaired two-tailed t test) and its rhythm was blunted across the light-dark cycle (Figure 1G). In addition, the analysis of the mRNA expression of the canonical clock-controlled gene *Dbp* shows that *Bmal1* deletion from MSNs induced a significant downregulation of this gene in the striatum (CTR versus SKO, P<0.005 for all timepoints, unpaired two-tailed t test) (Figure 1H). These results suggest that both *Per2* and *Dbp* expression in MSNs are controlled locally by *Bmal1* and that in the absence of *Bmal1* in the expression of these genes is disrupted.

### Sexually Dimorphic Effect of *Bmal1* Deletion from MSNs on Alcohol Intake and Preference

To study alcohol intake and preference, 12 - 18 weeks old male and female Bmal1CTR mice (males, n = 12; females, n = 17) and Bmal1SKO mice (males, n = 13; females, n = 14), were housed individually under a normal 12:12 h light-dark cycle, with food and water available *ad libitum*. For measurements of voluntary alcohol consumption, mice had free access to one bottle of 15 % ethanol solution in tap water (vl/vl) and one bottle of only tap water, every other day, in alternate left-right position, for a total of 11 sessions. Analysis of variance (ANOVA) revealed a significant main effect of genotype on alcohol intake (g/Kg/day) and preference (alcohol intake/total fluid intake) in both males (Figures 2A and B) and females (Figures 2C and D). Specifically, Bmal1SKO males consumed significantly more alcohol (P<0.002, ANOVA) and exhibited significantly greater alcohol preference (P<0.005, ANOVA) than Bmal1CTR mice over the 11 alcohol test days. On average, the daily alcohol intake (12.12 ± 0.64 g/Kg) and preference (0.80 ± 0.03) of Bmal1SKO male mice were 33 % and 36 % higher, respectively, than those of Bmal1CTR mice (average intake, 9.1 ± 0.62g/Kg; average preference, 0.62 ± 0.05; unpaired two-tailed t test, intake: P<0.003; preference: P<0.005, unpaired two-tailed t test). In contrast to males, Bmal1SKO females consumed significantly less alcohol than Bmal1CTR mice (P<0.03, ANOVA) and exhibited lower alcohol preference (P<0.05, ANOVA) over the 11 alcohol test days. The mean daily alcohol intake of Bmal1SKO females was 22 % lower than that of Bmal1CTR females (12.79 ± 1.29 versus 16.21 ± 0.88 g/Kg, P<0.03, unpaired two-tailed t test). The mean daily alcohol preference value of Bmal1SKO females was 15 % lower than that of Bmal1CTR females (0.60 ± 0.06 versus 0.75 ± 0.03, P<0.03, unpaired two-tailed t test).

**Figure 2.**
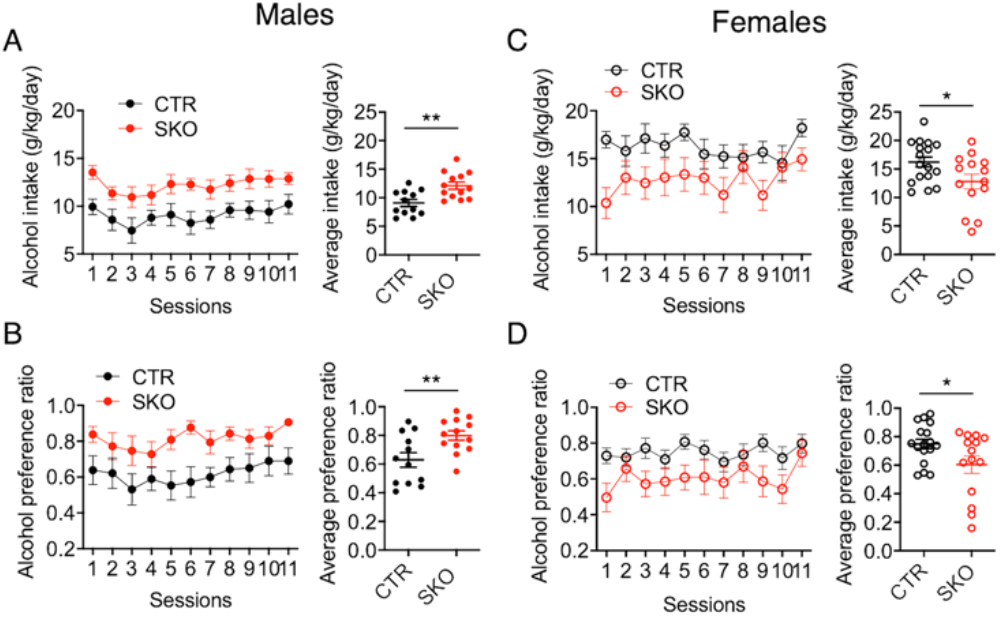
Opposite effects of striatal *Bmal1* deletion on alcohol intake and preference in male and female mice. **A)** Mean ± S.E.M. daily alcohol consumption (left) and average alcohol consumption (right) in control (CTR, n = 12) and *Bmal1* knockout (SKO, n = 13) male mice (**P<0.01, unpaired two-tailed t test). **B)** Mean ± S.E.M. daily alcohol preference (left) and average alcohol preference (right) in CTR and *Bmal1* SKO male mice (**P<0.01, unpaired two-tailed t test). **C)** Mean ± S.E.M. of daily alcohol intake (left) and average alcohol intake in CTR (n = 17) and Bmal1 SKO (n = 14) female mice (*P<0.05, unpaired two-tailed t test). **D)** Mean ± S.E.M. of daily alcohol preference (left) and average preference in CTR and Bmal1 SKO female mice (**P<0.05, unpaired two-tailed t test).

### Striatal Deletion of *Per2* Mimicks the Effect of *Bmal1* Deletion on Alcohol Intake in Males

*Bmal1* plays an obligatory role in transcriptional activation of the core clock gene *Per2*, and analysis of *Per2* mRNA in the striatum of Bmal1SKO and Bmal1HET mice revealed gene-dose dependent suppression of *Per2* expression at ZT5 and 17 (Figure 1G). *Per2* has been associated with alcohol consumption in humans, and global disruption of *Per2* has been shown to augment alcohol intake and preference in male mice [45], raising the possibility that the effect of striatal *Bmal1* deletion on alcohol intake and preference involves changes in local expression of *Per2*. MSNs that express D1 or D2 DA receptors and GPR88 also express PER2 (Figures 3A and 3B).

**Figure 3.**
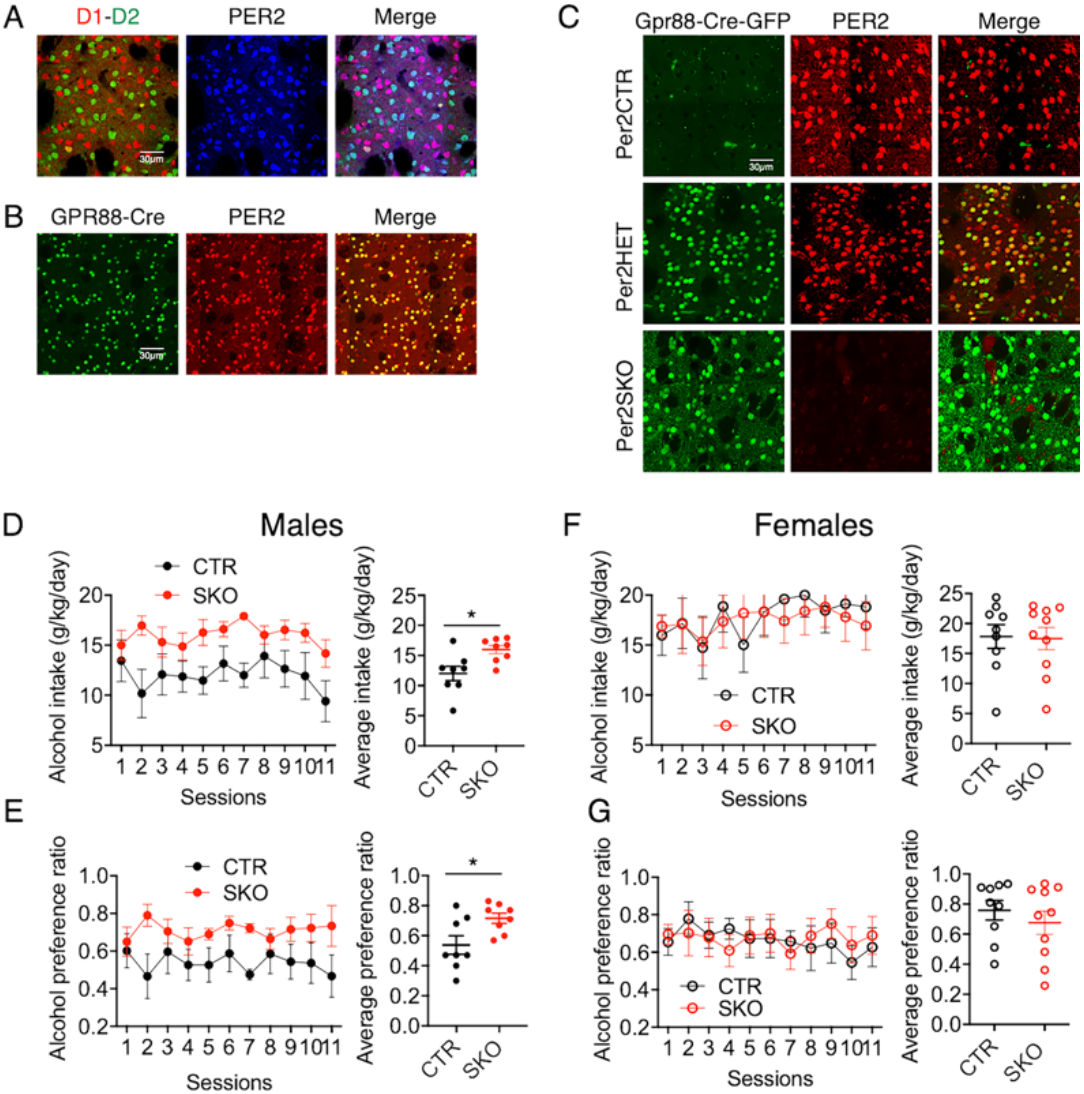
Alcohol consumption is altered in *Per2* knockout male mice. **A)** Representative image of immunofluorescence staining showing that PER2 (blue) is expressed in D1 (red) and D2 (green) dopamine receptor bearing neurons of the dorsal striatum in tdTomato-D1/GFP-D2 double transgenic reporter mice (scale bar = 30 μm). **B)** Representative image of immunofluorescence staining showing that PER2 (red) is expressed in Gpr88-Cre-GFP (green) positive MSN of the mouse dorsal striatum (scale bar = 30 μm). **C)** Representative image of immunofluorescence staining showing that *Per2* knockout mice which express Cre-GFP in dorsal striatal MSNs and have two floxed *Per2* alleles, display depleted PER2 in GFP-positive cells. *Per2* heterozygote mice which express Cre, but only possess a single floxed *Per2* allele, retain PER2 expression. Control mice which contain two floxed *Per2* alleles, but lack Cre, retain PER2 expression. **D)** Mean ± S.E.M. of daily alcohol consumption (left) and average alcohol consumption (right) in control (CTR, n = 8) and *Per2* knockout (SKO, n = 8) males (*P<0.05, unpaired two-tailed t test). **E)** Mean ± S.E.M. of daily alcohol preference (left) and average alcohol preference (right) in Per2CTR and Per2SKO male mice (*P< 0.05, unpaired two-tailed t test). **F)** Mean ± S.E.M of daily alcohol consumption (left) and average alcohol consumption (right) in Per2CTR (n = 9) and Per2SKO (n = 10) female mice. **G)** Mean ± S.E.M. of daily alcohol preference (left) and average alcohol preference (right) in Per2CTR and Per2SKO female mice.

To study the contribution of striatal *Per2* gene expression, we crossed C57BJ/6 mice homozygous for floxed alleles of the *Per2* locus (Per2fl/fl, European Mouse Mutant Archive, Strain ID: EM10599) with mice that express Cre and GFP under control of the *Gpr88* promoter to generate Per2SKO (Gpr88Cre/+; Per2fl/fl), Per2HET (Gpr88Cre/+; Per2fl/+) and Per2CTR (Gpr88+/+; Per2fl/fl) male and female mice. Immunostaining of striatal brain sections from Per2SKO mice revealed complete absence of PER2 immunoreactivity (Figure 3C). Deletion of *Per2* from MSNs augmented voluntary alcohol intake (Per2SKO versus Per2CTR, P<0.01, ANOVA) and preference (Per2SKO versus Per2CTR, P<0.05, ANOVA) in males (Figures 3D and E), thus mimicking the enhancing effect of striatal *Bmal1* deletion on male alcohol intake and preference. On average, the daily alcohol intake (15.99 ± 0.7g/Kg) and preference (0.71 ± 0.03) of Per2SKO male mice was ~33 % higher than intake (12.01 ± 1.2g/Kg) and preference (0.53 ± 0.06) in control littermates. In contrast, deletion of *Per2* from MSNs had no effect on alcohol consumption and preference in females (Figures 3F and G), revealing a female-specific dissociation between the effect of *Bmal1* and *Per2* on alcohol intake.

### Effect of Deleting a Single Copy of *Bmal1* or *Per2* from MSNs on Alcohol Intake

For a minority of genes, one functional copy is not sufficient to sustain normal function, and mutations causing the loss of function of one of the copies of such gene can impact behavior [57]. To determine if alcohol drinking behavior is affected when only one copy of *Bmal1* is expressed, we assessed alcohol intake and preference in male and female Bmal1HET mice as described above. Although not statistically significant, Bmal1HET males tended to consume and prefer more alcohol than Bmal1CTR male mice (Figure 4A and B). In contrast, Bmal1HET females consumed significantly less alcohol (P<0.02, ANOVA) and had lower preference (P<0.02, ANOVA) than Bmal1CTR females (Figure 4C and D). These results indicate that the *Bmal1* gene in the MSNs of the striatum is haploinsufficient regarding normal alcohol consumption and preference in females, whereas in males, deletion of one copy of striatal *Bmal1* appears not to affect alcohol intake and preference. Deletion of one copy of *Per2* (Per2HET) had no effect on alcohol intake and preference in males or in females, indicating that striatal *Per2* is haplosufficient for alcohol intake (Figures 4E, F, G, and H).

**Figure 4.**
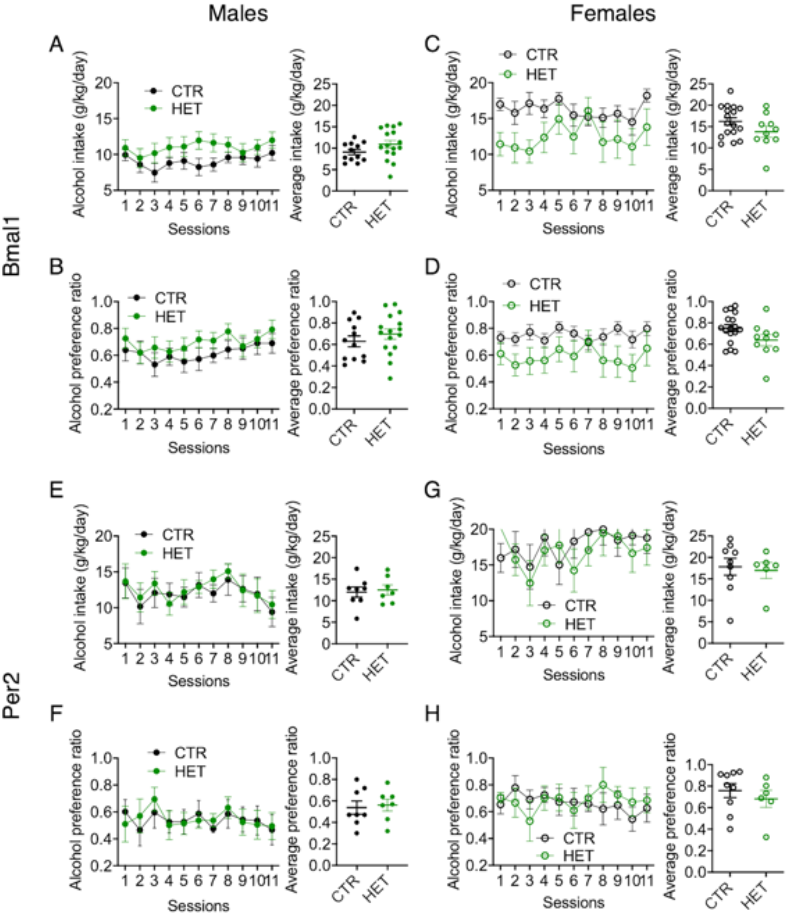
Alcohol consumption is altered in *Bmal1* heterozygote female mice. **A)** Mean ± S.E.M. of daily alcohol consumption (left) and of average consumption (right) in Bmal1CTR (n = 12) and Bmal1HET (n = 16) male mice. **B)** Mean ± S.E.M. of daily alcohol preference (left) and of average preference (right) in Bmal1CTR and Bmal1HET male mice **C)** Mean ± S.E.M. of daily alcohol consumption (left) and average consumption (right) in Bmal1CTR (n = 17) and Bmal1HET (n = 10) female mice. **D)** Mean ± S.E.M. of daily alcohol preference (left) and average preference (right) in Bmal1CTR and Bmal1HET female mice. **E)** Mean ± S.E.M. of daily alcohol consumption (left) and average consumption (right) in Per2CTR (n = 8) and in Per2HET (n = 7) male mice. **F)** Mean ± S.E.M. of daily alcohol preference (left) and average preference (right) in Per2CTR (n = 8) and in Per2HET (n = 7) male mice. **G)** Mean ± S.E.M. of daily alcohol consumption (left) and average consumption (right) in Per2CTR (n = 9) and in Per2FET (n = 6) female mice. **H)** Mean ± S.E.M. of daily alcohol preference (left) and average preference (right) in Per2CTR and in Per2FET female mice.

### *Gpr88* Monoallelic Expression Does Not Affect Alcohol Intake

Bmal1SKO and Per2SKO mice and their respective HET counterparts have only one functional copy of *Gpr88* in MSNs since one copy is modified to drive *Cre* and GFP expression. Complete deletion of *Gpr88* has been shown to augment alcohol intake in male mice [58, 59], raising the possibility that the changes in alcohol consumption seen in striatal Bmal1SKO and striatal Per2SKO were due, at least in part, to a sex difference in a contributory effect of *Gpr88* monoallelic expression. To study this possibility, we compared alcohol intake and preference between Gpr88+/+ mice and Gpr88Cre/+ mice. As shown in Supplementary Figure 1A and C, average alcohol intake and preference in Gpr88Cre/+ males were similar to those in Gpr88+/+ control male mice. Similarly, average alcohol intake and preference in Gpr88Cre/+ females were similar to those in Gpr88+/+ control female mice (Supplementary Figures 1B and D). These results exclude the possibility that the changes in alcohol consumption associated with deletion of striatal *Bmal1* or *Per2* were due to *Gpr88* haploinsufficiency, or expression of Cre-EGFP in Gpr88 bearing cells.

### Striatal deletion of Bmal1 or Per2 do not affect total fluid intake or sucrose consumption

Contrary to the effects of *Bmal1* deletion on alcohol intake and preference, deletion of *Bmal1* had no effect on total daily fluid intake across the 11 test sessions in either males or females (supplementary Figures 2A and B). Similarly, in both males and females, deletion of *Per2* had no effect on total fluid intake (Supplementary Figures 2C and D). Moreover, striatal deletion of *Bmal1* or *Per2* had no effect on voluntary intake of 0.25% or 2.0% sucrose solution (each solution was given in a two-bottle choice with water for 3 consecutive days) in males (Supplementary Figures 2E and G) or females (Supplementary Figures 2F and H). Thus, the effects of striatal deletion of *Bmal1* or *Per2* on alcohol intake and preference in both male and female mice appear not to be the result of global effect on either fluid consumption, general reward processing, or changes in caloric needs.

### Striatal deletion of *Bmal1* or *Per2* Do Not Affect Circadian Wheel-Running Behavior

Global deletion of *Bmal1* or *Per2* or selective deletion from the SCN disrupts circadian behavioral rhythms in animals [5, 60], and disruption of circadian rhythms can influence alcohol consumption [61–63]. To determine whether the effects of *Bmal1* or *Per2* deletion in the striatum on alcohol intake and preference was not related to changes in circadian rhythms induced by ectopic Cre recombinase expression, we monitored wheel-running behavior in alcohol naïve mice (n = 7-9 mice/genotype/sex) housed individually under different lighting conditions. The representative actograms shown in Supplementary Figure 3A show that circadian wheel running rhythms in Bmal1SKO and Bmal1HET male and female mice were indistinguishable from those in respective male and female Bmal1CTR mice. Similarly, wheel running rhythms in Per2SKO and Per2HET male and female mice were indistinguishable from those in respective Per2CTR mice (Supplementary Figure 3B). In particular, *Bmal1* or *Per2* deletion did not affect entrainment of daily wheel running to a 12:12 h light-dark cycle, adjustments to phase delay and advance shifts in the light cycle, and free running in constant conditions. These results show that Bmal1SKO and Per2SKO male and female mice and their respective HET mice have a functional SCN molecular clock and normal circadian pacemaking. Thus, the changes in alcohol consumption in SKO and HET male and female mice are independent of the SCN clock and not the result of disrupted circadian behavioral rhythms.

### Deletion of *Bmal1* or *Per2* from MSNs Eliminates Sex Differences in Alcohol Consumption

Sex differences in alcohol intake and preference are well recognized in animals and humans, with females generally drink more alcohol than males and are at a greater risk of developing alcohol dependence [27–29, 64–66]. Consistent with this, average daily alcohol intake in female Bmal1CTR mice was significantly greater than in Bmal1CTR males (P<0.0001, unpaired two-tailed t test) and mean alcohol preference was significantly higher in Bmal1CTR females than Bmal1CTR males (P<0.05, unpaired two-tailed t test), which corresponds to a 78 % greater intake and 21 % higher preference in females, respectively (Supplementary Figure 4A). In contrast, mean alcohol preference in Bmal1SKO male mice was 40 % greater and significantly different compared to Bmal1SKO females (P<0.005, unpaired two-tailed t test, Supplementary Figure 4C), and no difference was observed in mean alcohol intake between Bmal1SKO males and females (Supplementary Figure 4A). Similarly, average alcohol intake was 46 % greater in Per2CTR females compared to Per2CTR males (intake: P<0.001, unpaired two-tailed t test) and alcohol preference was 40 % higher in Per2CTR females than Per2CTR males (Supplementary Figures 4B and D). In contrast, there were no significant differences in mean alcohol preference and intake between Per2SKO males and females (Supplementary Figures 4B and D). These results show that selective deletion of *Bmal1* from MSNs eliminates sex differences in alcohol intake and preference by augmenting consumption and preference in males and suppressing the heightened intake and preference in females. Likewise, striatal deletion of Per2 eliminates sex differences in alcohol consumption, although the effect is attributed primarily to augmentation of intake and preference in Bmal1SKO males.

## Conclusions

Polymorphisms in *Bmal1* have been associated with alcohol consumption and alcohol use disorder. Our work reveals, for the first time, a causal role of *Bmal1* in striatal control of alcohol consumption. The influence of striatal *Bmal1* is sexually dimorphic, associated with repression of alcohol preference and intake in male mice, possibly via effect of *Per2*, and with enhancement of preference and intake in females via a mechanism independent of *Per2*. Strikingly, while wild-type mice exhibit the well-established heightened female-specific alcohol intake and preference, the opposing effects of *Bmal1* deletion on alcohol intake in males and females abolish this difference. This suggests that sexual dimorphism in the effect of *Bmal1* in the striatum contributes to sex differences in alcohol consumption in mice. The mechanisms that mediate this sexually dimorphic influence are as yet unknown but could involve sex-specific interactions of *Bmal1* with sex hormone receptors and dopamine signaling in MSNs [32, 67–77]. Transcriptomic analysis of differential striatal gene expression in this mouse model could define the downstream outputs of clock gene expression that affect sex differences in alcohol consumption. In summary, our findings uncover a novel *Bmal1*-linked mechanism in striatal MSNs that modulates alcohol intake in a sexually dimorphic manner, and which may contribute to sex differences in alcohol consumption in mice. Whether a similar mechanism in the striatum or elsewhere in the brain contributes to sex differences in other behaviors and disorders remains to be investigated in order to gain better insight into the sexually dimorphic influence of *Bmal1* on behavior.

## Supporting information

BIORXIV/2020/299842

## Supplemental Information

Supplemental Information includes four figures and Supplemental Experimental Procedures.

## Acknowledgments

This work was funded by grants from the Canadian Institutes of Health Research (S.A). We thank M. Parent, Laval University for the generous gift of tdTomato-D1/GFP-D2 double transgenic mice.

## Notes

### Competing Interest Statement

The authors have declared no competing interest.

