## Supplementary material for "Sexually dimorphic influence of the circadian clock gene *Bmal1* in the striatum on alcohol intake": BIORXIV/2020/299842

**Supplementary Fig. 1**

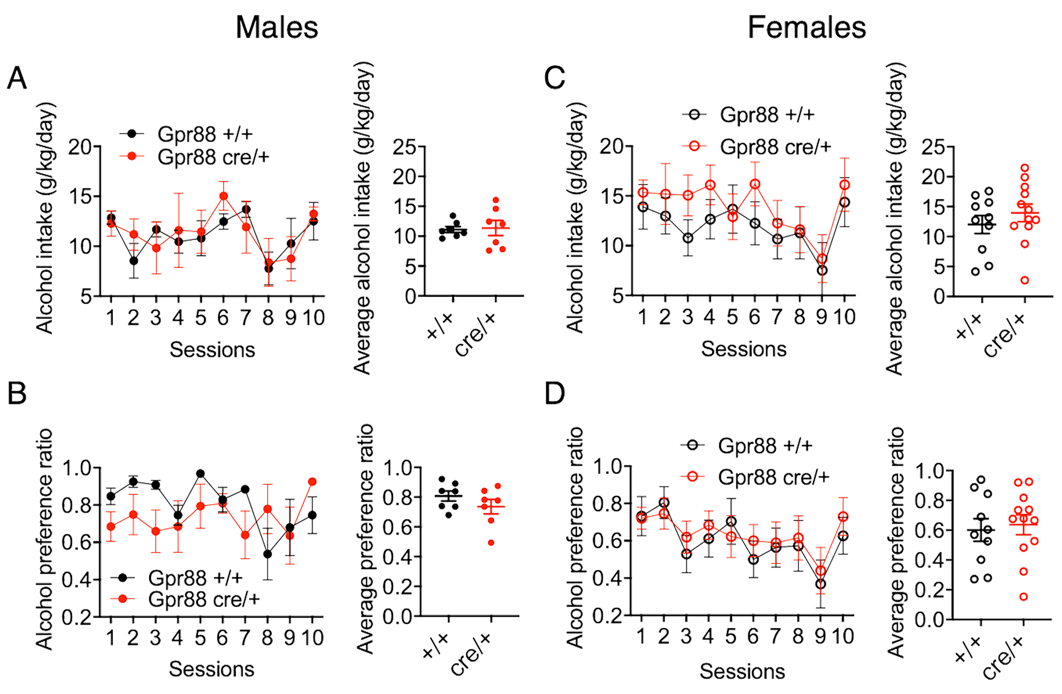

**A)** No difference in daily alcohol consumption (left) and average alcohol consumption (right) between male Gpr88+/+ (n = 7) and male Gpr88Cre/+ (n = 7) mice.

**B**) No difference in daily alcohol preference (left) and average alcohol preference (right) between male Gpr88+/+ and male Gpr88Cre/+ mice.

**C)** No difference in daily alcohol consumption (left) and average alcohol consumption (right) between female Gpr88+/+ (n = 10) and female Gpr88Cre/+ (n = 12) mice.

**D**) No difference in daily alcohol preference (left) and average alcohol preference (right) between female Gpr88+/+ and female Gpr88Cre/+ mice.

**Supplementary Fig. 2**

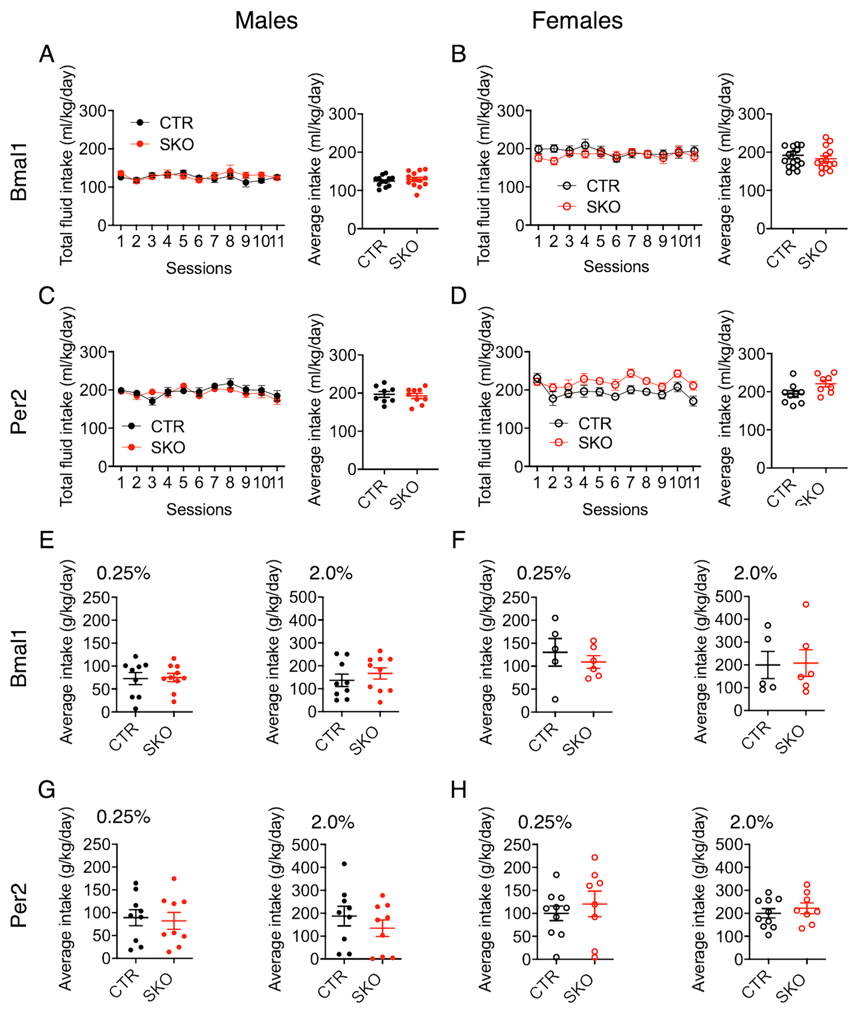

**A**) No difference in daily fluid intake (left) and average fluid intake (right) between Bmal1CTR and Bmal1SKO male mice.

**B**) No difference in daily fluid intake (left) and average fluid intake (right) between Bmal1CTR and Bmal1SKO female mice.

**C**) No difference in daily fluid intake (left) and average fluid intake (right) between Per2CTR and Per2SKO male mice.

**D**) No difference in daily fluid intake (left) and average fluid intake (right) between Per2CTR and Per2SKO female mice.

**E)** No difference in average daily intake of 0.25 % sucrose solution (left) or 2 % sucrose solution (right) between male control (n = 9) and Bmal1SKO mice (n = 10).

**F**) No difference in average daily intake of 0.25 % sucrose solution (left) or 2 % sucrose solution (right) between Bmal1CTR (n = 5) and Bmal1SKO (n = 6) female mice.

**G)** No difference in average daily intake of 0.25 % sucrose solution (left) or 2 % sucrose solution (right) between Per2CTR (n = 9) and Per2SKO (n = 9) males.

**D**) No difference in average daily intake of 0.25 % sucrose solution (left) or 2 % sucrose solution (right) between Per2CTR (n = 10) and Per2SKO (n = 8) females.

**Supplementary Fig. 3**

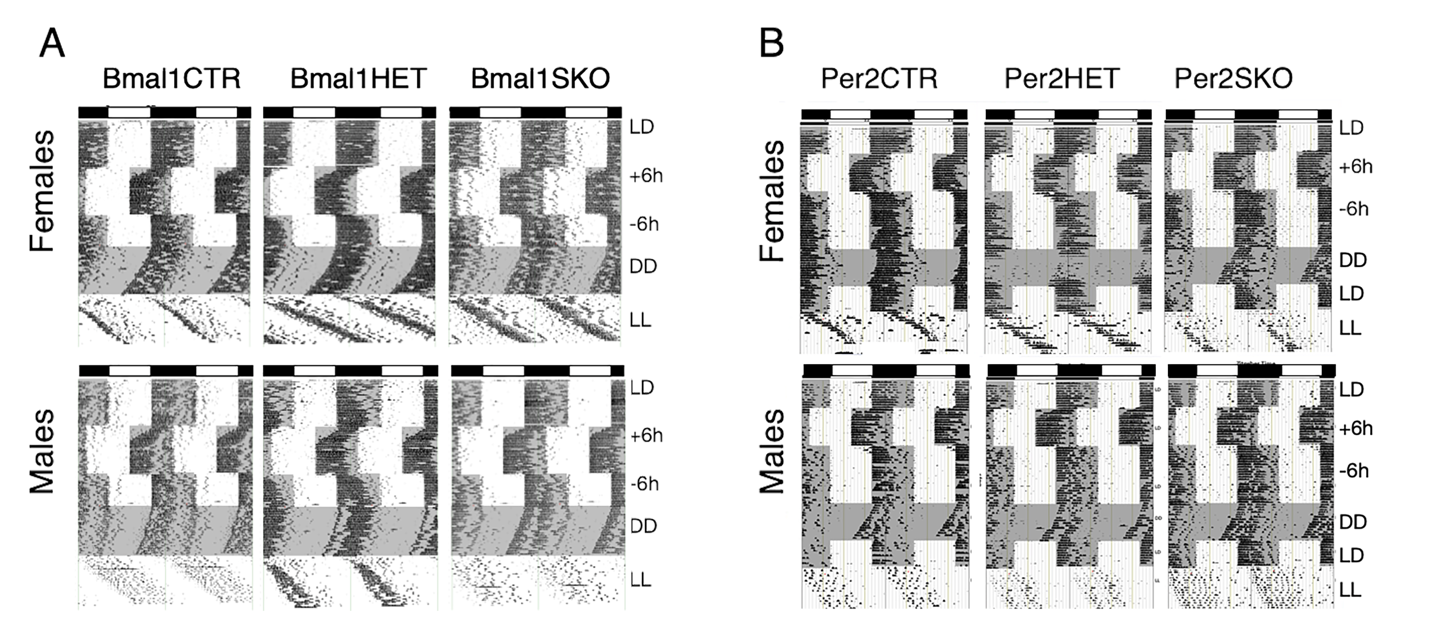

**A)** Representative double-plotted actograms illustrating the daily pattern of running-wheel activity in Bmal1CTR, Bmal1HET and Bmal1SKO female (top) and male (bottom) mice.

**B)** Representative double-plotted actograms illustrating the daily pattern of running-wheel activity in Per2CTR, Per2HET and Per2SKO female (top) and male (bottom) mice.

The vertical marks indicate periods of activity of at least 10-wheel revolutions per 10 min. Each horizontal line plots 48 h and sequential days are arranged from top to bottom. The light and dark phases are illustrated by the empty and gray shaded areas in each actogram, respectively (LD, 12:12 h light-dark; +6h, 6h phase advance; -6h, 6h phase delay; DD, constant dark; LL, constant light).

**Supplementary Fig. 4**

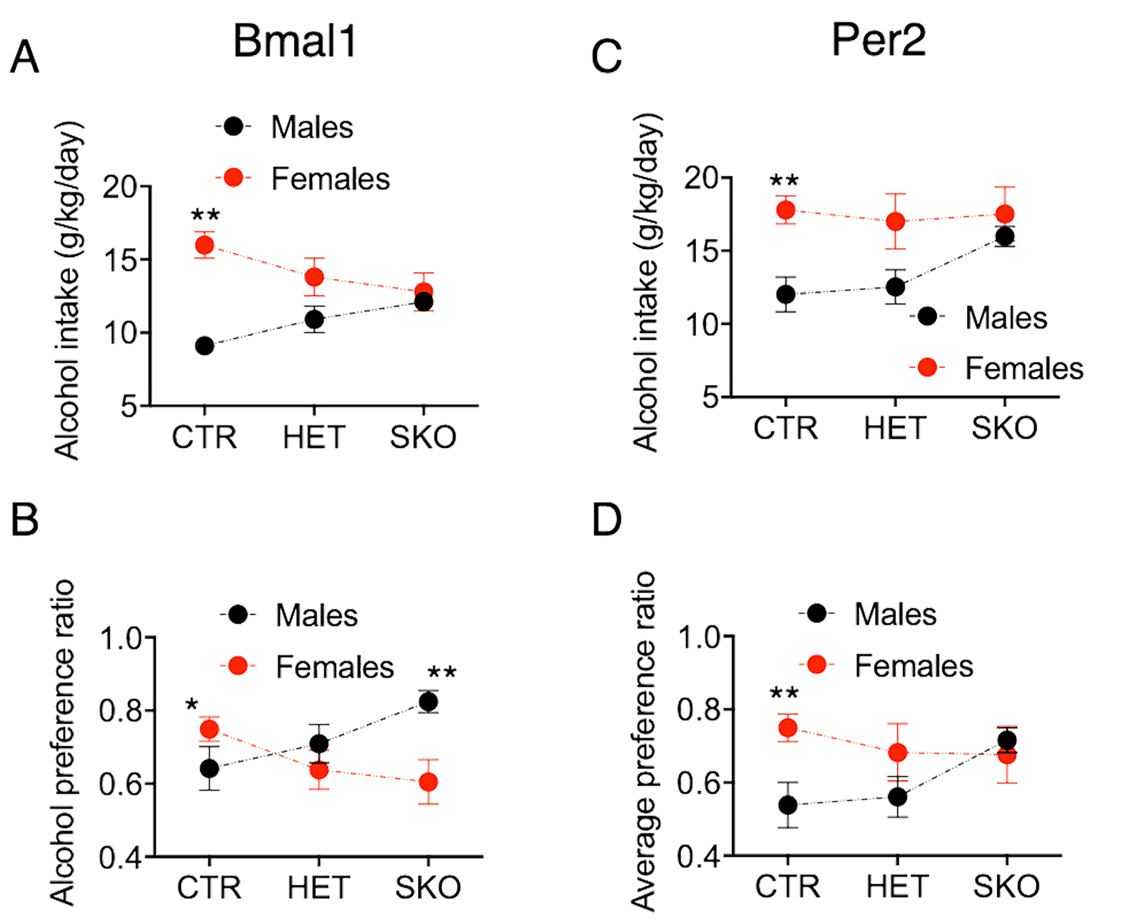

A) Average alcohol consumption in Bmal1CTR, Bmal1HET and Bmal1SKO males (n = 12 CTRL, 16 HET, 13 SKO) and female (n = 17 CTR, 10 HET, 14 SKO) mice (**P<0.001, unpaired two-tailed t test)

B) Average alcohol preference in Bmal1CTR, Bmal1HET and Bmal1SKO males

and female mice (*P<0.05, **P<0.001, unpaired two-tailed t test)

C) Average alcohol consumption in Per2CTR, Per2HET and Per2SKO males (n = 8 CTR, 7 HET, 8 SKO) and female (n = 9 CTR, 6 HET, 10 SKO) mice (**P<0.001, unpaired two-tailed t test).

D) Average alcohol preference in Per2CTR, Per2HET and Per2SKO males

and female mice (**P<0.001, unpaired two-tailed t test).

**Supplemental Experimental Procedures**

**Animals and genotyping**

Conditional knockout mice lacking BMAL1 or PER2 protein in the striatum were generated by two genetic crosses. In a first cross, *Gpr88*(Cre/+) male mice (B6.129S4-*Gpr88^tm1.1(cre/GFP)Rpa^*/J; stock number 022510; Jackson Laboratory) were bred with *Bmal1*(fl/fl) (B6.129S4(Cg)-*Arntl^tm1Weit^*/J; stock number 007668; Jackson Laboratory) or *Per2*(fl/fl) (B6.129-*Per2^tm1.2Ual^*/Biat, strain ID: EM10599, European Mouse Mutant Archive) female mice to generate respective heterozygote F1 progeny ([Gpr88Cre/+; Bmal1fl/+] or [Gpr88Cre/+; Per2fl/+]). In a second step, F1 males were crossed with *Bmal1*(fl/fl) or *Per2*(fl/fl) females to generate desired experimental and control animals. All floxed and *Cre*-expressing transgenic mouse lines have been backcrossed onto a C57BL/6J background for at least 6 generations. Genomic DNA was isolated from tail biopsies (3), and genotyping was performed by polymerase chain reaction (PCR) using the OneTaq® Hot Start 2X Master Mix (M0484, New England Biolabs Inc., Ipswich, MA, USA) according to the manufacturer’s instructions. The specific primers for genotyping were: *Cre*: Fwd-5’-TTTTCACCTCCCTCCCTTCT-3’; Rev-5’- GCCCACGATTCTTCTTCCTC-3’, yielding a 255 bp product for wild type or a 180 bp product for mutant mice; *Bmal1*: Fwd-5’-CTGGAAGTAACTTTATCAAACTG-3’, Rev-5’-CTGACCAACTTGCTAACAATTA-3’, yielding a 327 bp product for wild type or a 431 bp product for mutant animals; *Per2*: Fwd-5’-CTGTGTCCCTGGTTTCTG-3’, Rev-5’-GCAGGGCAGTTTCATCAAGG-3’, yielding a 468 bp product for wild type or a 596 bp product for mice carrying the transgene. tdTomato-D1/GFP-D2 double transgenic mice, provided by M. Parent, Laval University were genotyped by the same method as described above using the following primers: *D1*^tdTomato^: Fwd-5′-CTTCTGAGGCGGAAAGAACC-3′, Rev-5′-TTTCTGATTGAGAGCATTCG; *D2*-*GFP*: Fwd-5′-GAGGAAGCATGCCTTGAAAA-3′, Rev-5′-TGGTGCAGATGAACTTCAGG-3′; yielding a 300 bp and 600 bp product for transgenic animals, respectively. Mice were group-housed (2-4 individuals) under a 12:12 hour light-dark cycle (08:00 am to 08:00 pm) with controlled temperature (22 ± 2°C) and humidity (40-60 % relative humidity), and had access to water and standard rodent chow (5057, Charles River Laboratories, Wilmington, MA, USA) *ad libitum*. Conditional *Bmal1* and *Per2* knockout mice (Gpr88Cre/+; Bmal1fl/fl [Bmal1SKO], Gpr88Cre/+; Per2fl/fl [Per2SKO]), littermate heterozygote (Gpr88Cre/+; Bmal1fl/+ [Bmal1HET], Gpr88Cre/+; Per2fl/+ [Per2HET]) and wild type control animals (Gpr88+/+; Bmal1fl/fl [Bmal1CTR], Gpr88+/+; Per2fl/fl [Per2CTR]) of both sexes as well as Gpr88Cre/+ and corresponding control male and female mice were used for experiments with an age of 12-18 weeks. All animal experimental and animal care procedures in this report were conducted in accordance with the guidelines set forth by the Canadian Council of Animal Care and by the Animal Care Committee of Concordia University.

**Quantitative PCR**

Brains were isolated and flash-frozen from 11 to 13-week-old female Bmal1SKO, HET and CTL mice at ZT5, ZT11, or ZT17 (5, 11 and 17 hours after lights turned on). 100 μm coronal section were obtained using a Microm HM 505 E Cryostat (Microm International, Walldorf, Germany) to collect brain tissue. RNA was extracted from tissue punches of the dorsal striatum (DS) and nucleus accumbens (NAcc) using the RNeasy Lipid Tissue Mini Kit (Qiagen, Hilden, Germany) and converted into cDNA by the iScript™ cDNA Synthesis Kit (Bio-Rad, Hercules, CA, USA) according to the manufacturer’s instructions. qPCR was performed using the following primer pairs: *mBmal1*: Fwd-5’-GCAGTGCCACTGACTACCAAGA-3’, Rev-5’-TCCTGGACATTGCATTGCAT-3’; *mPer2:* Fwd-5’-TGATCGAGACGCCTGTGCTCG-3’, Rev-5’-CTCCACGGGTTGATGAAGCTG-3’; *mDbp*: Fwd-5’-AAGAAGGCAAGGAAAGTCCAG-3’, Rev-5’-ACCTCTTGGCTGCTTCATTG-3’; *mGapdh*: Fwd-5’-AGGTCGGTGTGAACGGATTTG-3’, Rev-5’-TGTAGACCATGTAGTTGAGGTC-3’. EvaGreen based amplification detection (EVOlution master mix, MBI-E250, Montreal Biotech, Canada) was performed using an Eco™ real-time PCR system (Illumina, San Diego, CA, USA). Relative expression of *Bmal1*, *Per2* and *Dbp* in relation to *Gapdh* was calculated based on the “delta delta C_t_” (ΔΔC_t_) method (4) and normalized to the highest expression value across genotypes and time points for each gene.

**Western blot analysis**

Mice were adapted for 10 d to a 12:12 h light-dark cycle and sacrificed either 2 hours (ZT2) or 14 hours (ZT14) after lights turned on. Brains were isolated and flash-frozen in isopentane. 200 μm coronal sections were cut on a cryostat (Microm HM 505 E Cryostat, Microm International, Walldorf, Germany). The striatal tissue was excised using punches and frozen on dry ice. Tissue was then homogenized in 150 μl of lysis buffer [1M Tris-HCl pH 6.8, 10 % sodium dodecyl sulfate, 0.1 ml phosphatase inhibitor cocktail 2 (Sigma, # P5726, Burlington, MA, USA), 0.1 ml phosphatase inhibitor cocktail 3 (Millipore Sigma, P0044, Burlington, MA, USA), 1× protease inhibitor cocktail (Roche, Basel, Switzerland)]. Protein extracts (40 μg/lane) were electrophoresed into a 10% SDS-PAGE gel and then transblotted onto a nitrocellulose membrane (Bio-Rad, # 1620112, Hercules, CA, USA). Membranes were blocked in 5% Milk TBST buffer (skim milk powder) and then incubated (overnight, 4°C) in TBST (with 5 % skim milk powder) with the anti-Bmal1 (1:1000 dilution) antibody (Novus Biologicals, # NB100-2288, Littleton, CO, USA). The same antibody was used for immunolabeling analysis. Next, the membrane was incubated in TBST (with 5 % milk) with a goat anti-rabbit IgG horseradish peroxidase-conjugated antibodies (1:200 dilution; Millipore Sigma, # AP132P, Burlington, MA, USA). The signal was visualized using the Western Lighting Chemiluminescence light-emitting system (PerkinElmer Life Sciences, Waltham, MA, USA). Between each antibody treatment, membranes were washed a minimum of three times (10 min per wash) in TBST.

**Tissue preparation for Immunohistochemistry**

Mice were transcardially perfused by cold saline (0.9 % NaCl), followed by cold paraformaldehyde solution (4 % in a 0.1 M phosphate buffer, pH 7.3). Brains were extracted and stored overnight in 4 % paraformaldehyde solution at 4 °C. Coronal sections were collected using a Leica vibratome, sliced at a thickness of 50 μm (immunohistochemistry) or 30 μm (immunofluorescence) and then stored at -20 °C in Watson’s cryoprotectant [35] until further use.

**Immunohistochemistry**

Free-floating sections were rinsed once for 10 min in phosphate buffered saline (PBS, pH 7.4), followed by 3 x 10 min rinses in 0.3 % Triton-X in Trizma-buffered saline solution (TBST: 0.3 % Triton, 50 mM Trizma buffer, 0.9 % saline). Immunohistochemistry for BMAL1 was performed using an affinity-purified rabbit polyclonal antibody, raised against BMAL1 (1:10000 Novus Biologicals # NB100-2288, Littleton, CO, USA). Brain sections were incubated (40 h, 4 °C) in a primary solution with Bmal1 polyclonal rabbit antibody, 2 % normal goat serum, 5 % milk buffer in TBST. The sections were then incubated in a secondary solution, composed of biotinylated anti-rabbit IgG, raised in goat (1:200, Vector Laboratories, Burlington, ON, Canada). Lastly, the sections were incubated in a tertiary Avidin-Biotin-Peroxidase solution (Vectastain Elite ABC Kit, Vector Laboratories, Burlington, ON, Canada). All sections were rinsed in a 0.5 % 3,3-diaminobenzidine (DAB) solution. Immunoreactive (IR) cells were stained using a 0.5 % DAB, 0.01 % H_2_O_2_ and 8 % NiCl_2_ solution. Stained sections were mounted onto gel-coated slides, dehydrated in a series of alcohols and Citrisolv (Fisher Scientific, Pittsburgh, PA, USA), covered by Permount media (Fisher Scientific, Pittsburgh, PA, USA) and coverslipped. The sections were examined under a light microscope (Leica, DMR) and identified using Paxinos mouse brain atlas [1]. Images of the SCN, Striatum and hippocampus were captured using a Sony XC-77 video camera, Scion LG-3 frame grabber (Scion Corporation, Frederick, MD, USA), and Image J [2].

**Immunofluorescence**

Free-floating sections were rinsed once for 10 min in phosphate buffered saline (PBS, pH 7.4), followed by 3 x 10 min rinses in 0.3 % Triton-X in PBS (PBST). Tissue was pre-blocked for 1 h at room temperature with gentle agitation in a solution of PBST containing 3 % skim milk powder and 6 % normal donkey serum (NDS) then directly transferred to the primary antibody incubation. Tissue was incubated for 2 h with the primary antibody at room temperature with gentle agitation, rinsed 3 x 10 min in PBST, then incubated with the secondary antibody for 1 h at room temperature with gentle agitation. Antibodies were diluted in a solution of 0.3 % PBST with 3 % skim milk powder and 2 % NDS. The following antibodies and dilutions were used: PER2 rabbit polyclonal (1:500, Novus Biologicals, # NB300-564, Littleton, CO, USA), BMAL1 rabbit polyclonal (1:500, Novus Biologicals # NB100-2288, Littleton, CO, USA), anti-rabbit secondary Alexa-647 (1:500, Life Technologies, Carlsbad, CA, USA). Once all incubations were complete, the tissue was rinsed 3 x 10 min in PBST, and a another 10 min in PBS before being mounted onto slides, allowed to air dry and coverslipped with VECTASHIELD® Antifade Mounting Media (Vector Laboratories, Inc., Burlingame, CA, USA). Slides were left to cure overnight in the dark, sealed with clear nail polish and imaged over the next 5 days. Slides were stored in a slide box at 4 °C. Fluorescent images were captured using the 60x objective on an Olympus FV10i automated confocal laser scanning microscope at the Centre for Microscopy and Cell Imaging, Concordia University, Montreal, Canada. Brain regions of interest were determined using Paxinos mouse brain atlas [1]. All confocal parameters (pinhole, contrast, brightness, etc.) were held constant across all data sets from the same experiment.

**Two-bottle choice- intermittent access**

12 - 18 weeks old male and female mice (n = 6 - 17) were separated from group housing one week prior to the beginning of the experiment and single-housed under a 12:12 h light-dark cycle. Oral alcohol intake was determined using 24-h intermittent access to alcohol in a two-bottle choice drinking paradigm. Every other day, at ZT4 (4 hours after the lights were on), they were given 24 hours of concurrent access to one bottle containing 15 % ethanol (v/v) in tap water and one bottle containing water for 11 sessions of alcohol access. The bottles were weighed every day and the mice were weighed at the beginning and once a week during the experiment. The position (left or right) of each solution was alternated between sessions as a control for side preference. The possible loss of solutions due to the handling of the bottles, was controlled by weighing bottles in empty cages.

**Sucrose consumption**

12 - 18 weeks old male and female mice (n = 5 - 10) were single-housed under a 12:12 h light-dark cycle. Every day, at ZT4, they were given free access to one bottle containing 0.25 % sucrose in tap water (m/v) and 1 bottle containing tap water, in alternate left-right position. Sucrose solution was offered for 3 consecutive days. and the amount of fluid intake was recorded every day. One week later, the same procedure was performed with 2 % sucrose solution. The bottles were weighed every day and the mice were weighed at the beginning of the experiment.

**Locomotor activity rhythms**

12-15 weeks old male and female mice were housed individually in cages equipped with a running wheel with *ad libitum* access to food and water. Each cage was kept in a temperature-controlled, soundproof, ventilated isolation chamber and with computer-controlled light sources. Main light source was adjusted such that approximately 100-300 lux of light was equally distributed in the chamber. Mice were kept under a 12:12 h light-dark (LD) schedule, and thereafter exposed to 6 h phase advance, followed by a 6 h phase delay. After that, mice were transferred into constant darkness (DD) followed by exposure to constant light (LL). Mice were kept for 21-40 days under the respective light regimen. Wheel rotation was recorded using the VitalView program (Mini Mitter, Starr Life Sciences Corp., Oakmont, PA, USA) and double-plotted actograms were prepared using the Actiview software (Mini Mitter, Starr Life Sciences Corp., Oakmont, PA, USA).

**Statistical analysis**

For behavioral experiments, data were analyzed with (GraphPad Prism) unpaired two-tailed t test and two-way ANOVA with or without repeated measures (RM-ANOVA). Significant main effects and interactions of the ANOVAs were further investigated with the Bonferroni post hoc test.

1]. Franklin, Keith B. J., and George Paxinos. 1997. The mouse brain in stereotaxic coordinates.

2]. Rasband, W.S., ImageJ, U. S. National Institutes of Health, Bethesda, Maryland, USA, https://imagej.nih.gov/ij/, 1997-2018.

3]. Truett GE, Heeger P, Mynatt RL, Truett AA, Walker JA, Warman ML. Preparation of PCR-quality mouse genomic DNA with hot sodium hydroxide and tris (HotSHOT). Biotechniques. 2000;29(1):52-54. doi:10.2144/00291bm09

4]. Livak KJ, Schmittgen TD (2001) Analysis of relative gene expression data using realtime quantitative PCR and the 2∆∆C(T) Method. Methods 25(4): 402–408.
